# Enforced expression of phosphatidylinositol 4-phosphate 5-kinase homolog (PIPKH) alters phosphatidylinositol 4,5-bisphosphate distribution and the localization of small G-proteins

**DOI:** 10.1101/377465

**Authors:** Yanbo Yang, Miriam Park, Gregory D. Fairn

## Abstract

The generation of phosphatidylinositol 4,5-bisphosphate (PtdIns(4,5)P_2_) by phosphatidylinositol 4-phosphate 5-kinases (PIP5Ks) is essential for many of the functions including the control of cytoskeleton, signal transduction and endocytosis. Additionally, due to its presence in the plasma membrane and its anionic charge PtdIns(4,5)P_2_, together with phosphatidylserine, imbue the inner leaflet of the plasma membrane with a negative surface charge. This negative charge helps to define the identity of the plasma membrane as serves to recruit or regulate a multitude of proteins that contain polybasic domains or patches. Here we determine that the phosphatidylinositol 4-phosphate 5-kinase homolog (PIPKH) alters the subcellular distribution of PtdIns(4,5)P_2_ by re-localizing the PIP5Ks to endomembranes. Consistently, we find a redistribution of the PIP5K family members to endomembrane structures upon PIPKH overexpression that is accompanied by an accumulation of PtdIns(4,5)P_2_ and phosphatidylinositol 3,4,5-trisphosphate (PtdIns(3,4,5)P_3_), which further influences the distribution of endosomes and lysosomes. Additionally, we demonstrate that the accumulation of polyphosphoinositides increases their negative surface charge that in turn leads to the relocalization of surface charge probes as well as the polycationic proteins K-Ras and Rac1.

## Introduction

Phosphoinositides (PIPs), the phosphorylated derivatives of phosphatidylinositol, constitute a relatively minor fraction of the total cellular lipids^1^. For the most part, PIPs are synthesized locally on organelles by specific kinases while they are subject to degradation or consumption by a variety of phosphatases and phospholipase C isoforms^1^. The inositol ring has three accessible hydroxyl groups that by differential phosphorylation leads to the generation of seven PIP species^^1,2^^. Many of the individual PIPs display preferential accumulation on specific organelles and thus the presence of individual PIP species has been postulated to provide identity to its organelle^2^.

Another biophysical feature of organellar membranes that contributes to their identity is the surface charge of the cytosolic leaflets. Experimental evidence has demonstrated that the inner leaflet of the plasma membrane (PM) possesses the highest negative charge density of the organelles^3–5^. This membrane is not only rich in the anionic phospholipids phosphatidylserine and phosphatidylinositol but also PIPs including PtdIns(4,5)P_2_^6^. The negative surface charge of the inner plasma membrane together with PtdIns(4,5)P_2_ and PtdIns(3,4,5)P_3_ target numerous peripheral proteins to the PM via either electrostatic interactions or by serving as ligands for modular protein domains^7,8^. Indeed, many peripheral proteins target the PM via a coincidence sensing typically by possessing a lipid modification such as prenylation and a polybasic patch^9,10^.

Mammalian cells produce PtdIns(4,5)P_2_ using two related yet distinct mechanisms. The type I phosphatidylinositol 4-phosphate 5-kinases (PIP5Ks) are present throughout the eukaryotic kingdom and are responsible for the bulk of PtdIns(4,5)P_2_ synthesis^11,12^. The second pathway that synthesizes PtdIns(4,5)P_2_ is mediated by the type II phosphatidylinositol 5-phosphate 4-kinases that use PtdIns(5)P as a substrate^12^. The type I and II enzymes appear to have arisen from a common ancestor and display ≈30% sequence identity^13^. Despite this similarity the type I and type II kinases localize to different subcellular locations^6,14^. Previous results demonstrated that the plasmalemmal localization of the type I kinases are influenced by its substrate PtdIns(4)P and the anionic surface charge of the inner leaflet of the PM^6^. Additionally, a number of protein-protein interactions have been described for the type I kinases including the small G-proteins Arf6, Rho and Bruton’s Tyrosine Kinase^15–17^. The impact of protein-protein interactions on the activity and subcellular localization of the PIP5Ks is incompletely understood.

The phosphatidylinositol 4-phosphate kinase homolog (PIPKH) gene also referred to as PIP5K-like 1 (PIP5KL1) was identified based on sequence similarity to the other PIP5K isoforms^18^. Despite the importance of the PIP5Ks to cellular function, the role of PIPKH has not been obvious. Initial studies detected the mRNA transcript predominantly in the brain and testis of mice^18^. While more recent proteomic studies have found the protein in gastric epithelial cells^19^. PIPKH has the ability to interact with PIP5Ka and p to potentially regulate the production of PtdIns(4,5)P_2_^18^. Examination of normal and cancerous gastric cancer samples revealed that PIPKH expression is lost the majority (≈65%) of the samples^20^. Additionally, re-expression of PIPKH in the gastric cancer cell line BGC823 resulted in a decrease in both cell migration and proliferation^20^. How specifically PIPKH influenced these properties is unclear.

Despite these observations the function of PIPKH within the cell remains unknown. Bacterially expressed PIPKH does not possess kinase activity while immunoprecipitates of epitope-tagged PIPKH from human embryonic kidney 293T cells were found to exhibit low PIP5K activity^18^. However, site-directed mutagenesis of PIPKH suggested that it was not the source of the PIP5K activity^18^. Instead it was determined that immunoprecipitating PIPKH results in the co-capture of PIP5Ka and PIP5Kβ^18^. Taken together, the prevailing thought is that PIPKH is likely a pseudokinase with an unknown function. In this study we aimed to determine the impact of the expression of PIPKH in a cell otherwise lacking this protein.

## Results

### PIPKH recruits PIP5Ks to intracellular compartments and interacts with PIP5Ks in cells

We first examined the subcellular distribution of heterologously expressed human PIPKH in Chinese hamster ovary (CHO) cells. CHO cells transiently transfected with plasmids encoding mCherry (mCh)-tagged PIPKH were examined using spinning-disc confocal microscopy. As shown in Fig. 1a and Supplemental Figure 1, PIPKH localized to the PM as well as endocytic membranes that were stained by the endocytic tracer FM4-64. A previous study demonstrated that recombinant PIPKH does not possess the ability to bind to liposomes *in vitro*^18^. Thus, we suspected that the localization of the PIPKH is a result of protein-protein interactions. To date, the only known interacting partner for PIPKH is PIP5Kα and PIP5Kβ. Previously, we had demonstrated that the Type I PIP5Ks localize primarily to the PM in a macrophage cell line and we confirmed this observation using GFP or YFP-chimeras in CHO cells (Fig. 1b)^6^. This raises the possibility that the plasmalemmal pool of PIPKH was due to interactions with the α and β isoforms. Surprisingly, when co-expressed we found that PIPKH caused a relocalization of not only PIP5K α and β but also the γ isoform to internal membranes (Fig. 1b-c). Thus, enforced expression of PIPKH was able to over-ride the endogenous targeting determinants of the PIP5Ks.

**Figure 1.**
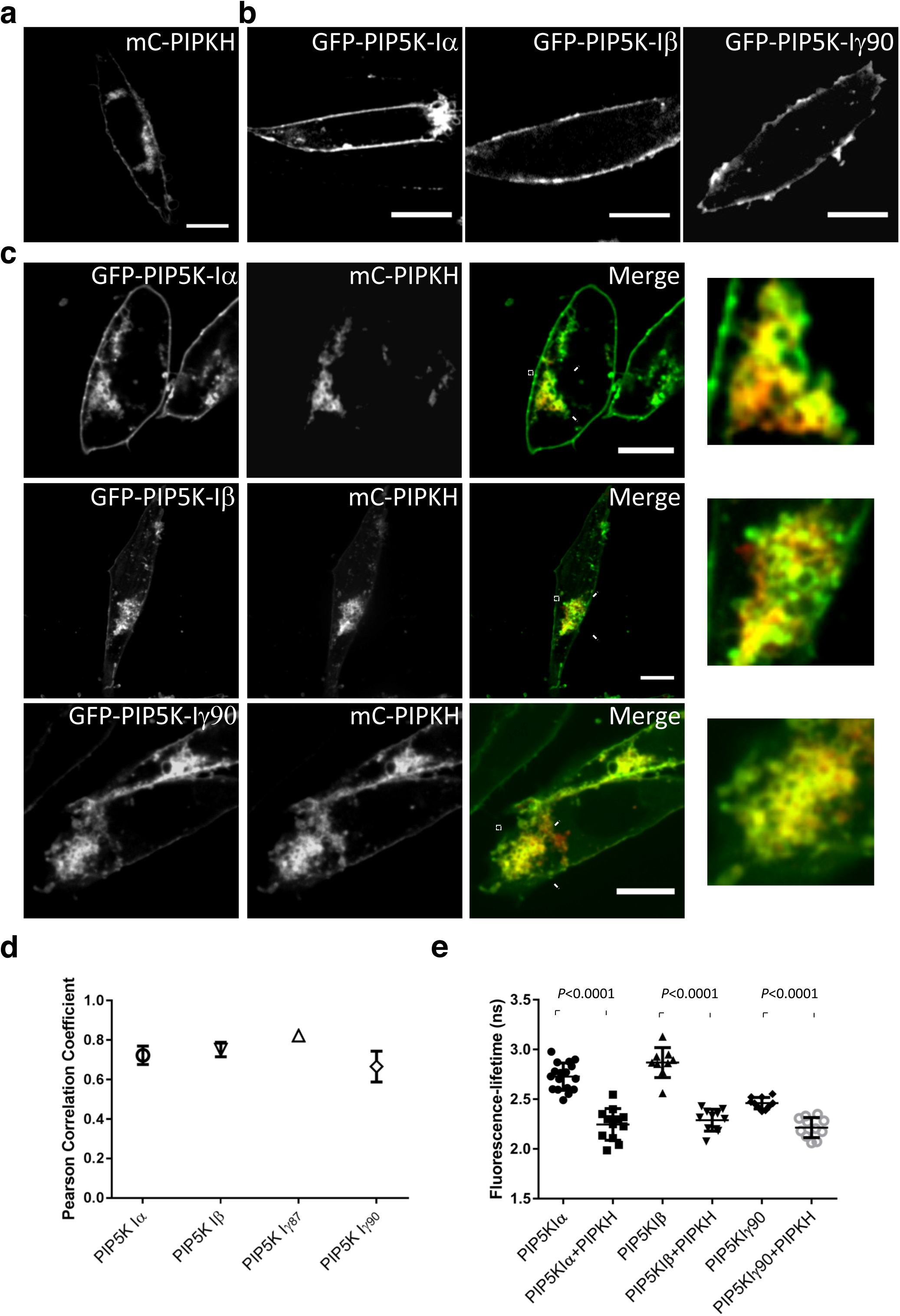
Enhanced expression of PIPKH redistributes PIP5Ks. (a) PIPKH localizes to the plasma membrane (PM) and endomembranes. Chinese Hamster Ovary (CHO) cells were transiently transfected with mCherry (mCh)-tagged PIPKH and observed by confocal microscopy. (b) Type I phosphatidylinositol 4-phosphate 5-kinases (PIP5Ks) localizes predominantly to the PM in CHO cells. Cells transiently transfected with GFP or YFP chimeras of PIP5K isoforms. (c) PIPKH over-expression causes a redistribution of PIP5Ks to internal structures. CHO cells co-transfected with the indicated YFP‐ or GFP-tagged PIP5K isoforms and mCh-tagged PIPKH. Images were acquired using spinning-disc confocal microscopy. Scale bars = 10 mm. (d) PIPKH and PIP5Ks extensively colocalize. Pearson’s correlation coefficient was determined for each of the indicated PIP5K isoforms and PIPKH. Values are the mean ± Std. Dev. With a minimum of 9 cells/condition imaged acquired from three separate experiments (n=3). (e) Fluorescently tagged versions of PIPKH and PIP5Ks are in close proximity to each other. Fluorescence Lifetime Imaging Microscopy – Forster Resonance Energy Transfer (FLIM-FRET) was used to examine the proximity of the GFP or YFP (donor) to the mCh-PIPKH (acceptor). Values present the mean lifetimes ±Std. dev. from a minimum of 9 cells observed in three separate experiments. Significance testing was performed using unpaired two-tailed Student’s t-test with Welch correction.

To determine if the PIPKH induced relocalization of the PIP5Ks was due to protein-protein interactions we used fluorescence-resonance-energy transfer (FRET) to detect the proximity of GFP/YFP-tagged PIP5K and mCh-PIPKH. More specifically, we took advantage of fluorescence lifetime imaging microscopy (FLIM) or so-called “FLIM-FRET” that obviates many of the technical issues associated with traditional FRET. Using FLIM-FRET, a decreased fluorescence lifetime of YFP-PIP5Kα/β and GFP-PIP5Kγ90 in cells that were transfected with mCh-PIPKH was observed (Fig. 1e). This result is consistent with the previous finding that demonstrated PIPKH and the α and β isoforms could co-immunoprecipitation together supporting the notion that they directly interact or could be part of a larger complex. These results suggest that PIPKH can heterodimerize with PIP5K isoforms and possibly act as a scaffold to localize and regulate the PIP5Ks.

### PIPKH causes intracellular accumulation of PtdIns(4,5)P_2_ and PtdIns(3,4,5)P_3_

As mentioned in the Introduction, a prevailing theory is that the presence of individual phosphoinositide species serves as molecular beckons to recruit peripheral proteins and provide identity to an organelle. As enforced expression of PIPKH caused the marked redistribution of PIP5Ks to endomembranes we wanted to consider if PIPKH could alter phosphoinositide distribution. First, we used several commonly used genetically encoded biosensors to monitor the distribution and relative abundance of individual PIP species. As depicted in Figure 2a, CHO cells have intracellular pools of PtdIns4P (as monitored by GFP-P4M), PtdIns3P (mCh-PX) and PtdIns(4,5)P_2_ (PH-PLC8) while these cells have little to no PI3,4P_2_ or PI3,4,5P_3_ (PH-AKT). Next, we contransfected these probes together with the PIPKH expressing plasmid. As depicted in Figure 2b, d the PIPKH displayed minimal over-lap with the monophosphorylated PIPs. Conversely, mCh-PIPKH caused a marked increase in both PI4,5P_2_ on the endomembranes (Fig2c). Additionally, these pools of PIPs co-localized very well with the PIPKH protein (Fig. 2c, d). These results suggest that the relocalized PIP5Ks remain active and that their substrate PI4P is rapidly converted to PtdIns4,5P_2_. Previous results had demonstrated that PIPKH does not impact the total cellular levels of PI4,5P_2_. With this in mind we examined determined the distribution of the PI4,5P_2_ probe in the plasma membrane vs. the rest of the cell. In control cells, nearly 100% of the PH-PLCδ is associated with the PM. However, in cells expressing PIPKH, ≈38% of the PI4,5P_2_ probe is now found in endomembranes.

**Figure 2.**
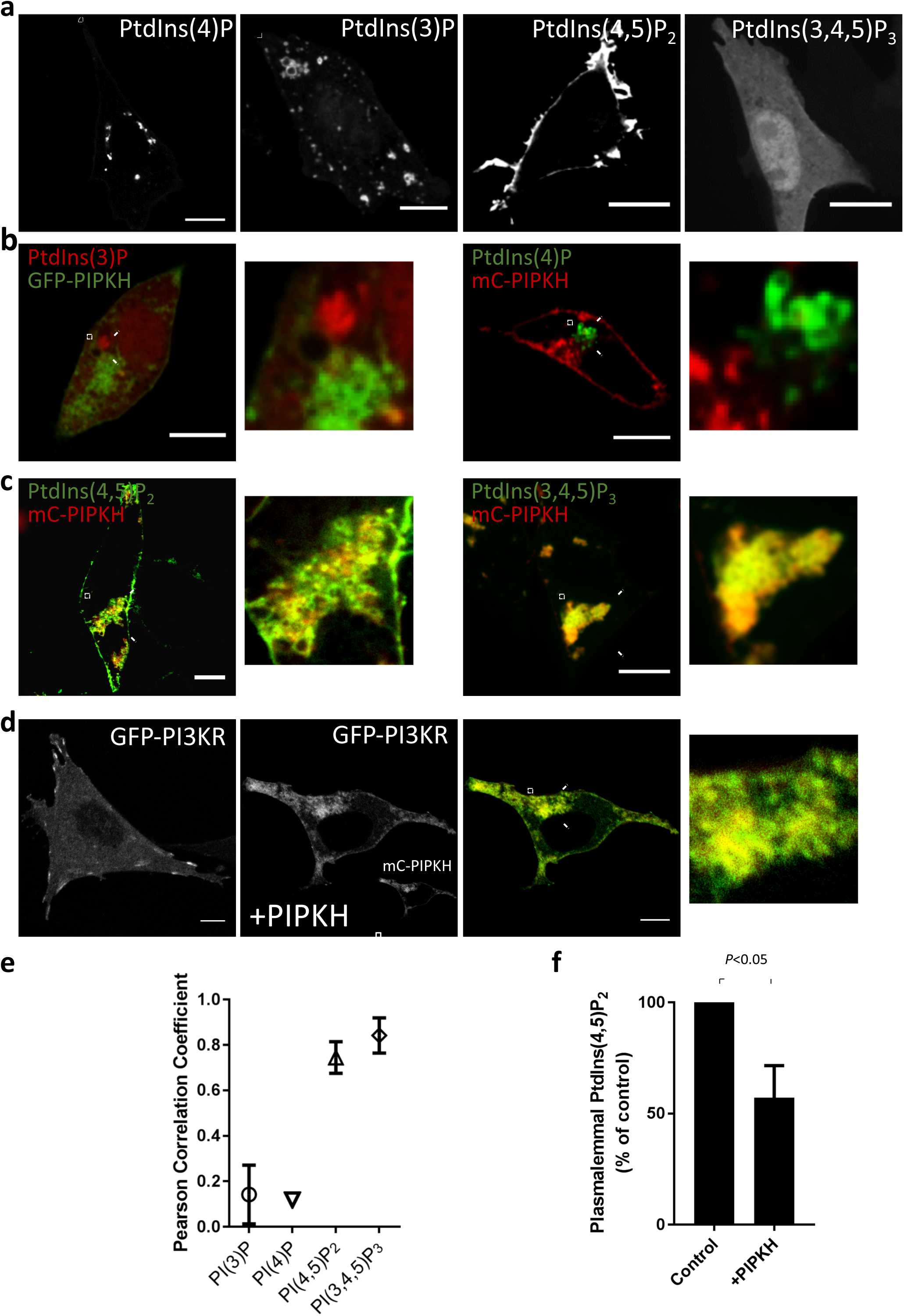
PIPKH results in an aberrant distribution of PtdIns(4,5)P_2_ and PtdIns(3,4,5)P_3_. (a) Distribution of phosphoinositides in CHO cells. CHO cells were transiently transfected with fluorescently-tagged probes for PtdIns(3)P (PX), PtdIns(4)P (tandem P4M), PI4,5P_2_ (PH-PLCδ), and PtdIns(3,4)Pz/PtdIns(3,4,5)P_3_ (PH-AKT). (b) PIPKH does not colocalize with mono-phosphorylated phosphatidylinositol. CHO cells were transiently co-transfected with mCh-PX and GFP-PIPKH or GFP-2xP4M and mCh-PIPKH were examined 18-24 h post-transfection. While both phosphoinositide probes decorate cytoplasmic structures, they are devoid of PIPKH. (c) PIPKH causes altered polyphosphoinositide distribution. CHO cells were transiently co-transfected with mCh-PIPKH and either GFP-PH-PLCδ or PH-AKT were examined 18 h post-transfection. PIPKH overlaps extensively with a cytoplasmic pool of PtdIns(4,5)P_2_ and PtdIns(3,4)P_2_/PtdIns(3,4,5)P_3_. Scale bars = 10 mm. (d) CHO cells transiently expressing PIPKH and the p85 subunit of phosphatidylinositol 3-kinase (PI3K) displayed a relocalization of the p85 from the plasma membrane to the endosomes. Scale bars = 10 mm. (e) Polyphohphoinositides but not mono colocalize with PIPKH. Pearson’s correlation coefficient of PIPKH with each of the indicated phosphoinositide probes. Values represent the mean ± Std. dev. from a minimum of 9 cells per experiment imaged from at least three different experiments (n=3). (f) Enforced expression alters the relative distribution of PtdIns(4,5)P_2_ in the cell. A histogram of the relative distribution of PtdIns(4,5)P_2_ in the cell as monitored by the PH-PLCδ probe. The data represents the percent of PH-PLCδ probe associated with the PM in control and PIPKH-overexpressing cells. The data represent the mean ± Std dev from a minimum of 9 cells per experiment from at least three different experiments (n=3). Significance testing was performed using unpaired two-tailed Student’s t-test with Welch correction.

Next, we sought to determine if this PtdIns(4,5)P_2_ could serve as a substrate for PtdIns(3,4,5)P_3_. Consistently, when co-expressed with mCh-PIPKH, GFP-tagged AKT-PH, a sensor for phosphatidylinositol 3,4-bisphosphate (PtdIns(3,4)P_2_) and PtdIns(3,4,5)P_3_, is co-localized with PIPKH to the cytoplasmic vesicular structures (Fig. 2c and d); similar results were also obtained using the PtdIns(3,4,5)P_3_ specific PH domain of BTK1 (data not shown). Consistent with these findings, we found that cells transiently expressing PIPKH and the p85 subunit of phosphatidylinositol 3-kinase (PI3K) displayed a relocalization of the p85 from the plasma membrane to the endosomes (Fig. 2d). Thus, transient overexpression of PIPKH results in the relocalization of at least two phosphoinositide kinases, PIP5Ks, and Class I PI3K, and thereby spatially alters the cellular distribution of PtdIns(4,5)P_2_ and PtdIns(3,4,5)P_3_.

### Localization of PIPKH on the limiting membrane of endosomes

The preceding results motivated us to determine which endosomal population(s) were being impacted by PIPKH. Importantly, as our experiments were impacting PtdIns(4,5)P2 we stained the cells with FM4-64 at 4°C to ensure that the internal PIPKH signal was detached from the PM and not in a long tubule (Supplemental Fig. 1). Next, we used GFP-chimeras of commonly used Rab proteins to identify specific endosomal compartments; Rab11a (recycling endosomes), Rab5 (early endosomes) and Rab7 (late endosomes/lysosomes) (Fig. 3a). In CHO cells the Rab11a-positive recycling endosomes tend to cluster near the microtubule organizing centre while the Rab5 and Rab7-positive populations have a more dispersed appearance. As illustrated in Fig. 3b-d, many of the PIPKH-induced clustered endosomes co-localized with Rab11a, Rab5, or Rab7. Transient transfection of fluorescently tagged Rab7 consistently gave us a high cytosolic or ER signal. To circumvent this issue, we compared the localization of the PIPKH with the Rab7-interacting lysosomal protein (RILP) a Rab7 effector that specifically binds to active GTP-bound form of Rab7, respectively (Fig. 3c)^21^. These results demonstrated that PIPKH extensively co-localized with active Rab7 along with the other Rab proteins. Finally, we assessed whether the PIPKH could be functionally interacting with the Rab proteins. To do this we again conducted the FLIM-FRET analysis and determined that the possibility of PIPKH interacting with Rab proteins is low (Fig. 3e). Collectively, these results suggest that PIPKH can localize to multiple types of endosomes and further influences their distribution.

**Figure 3.**
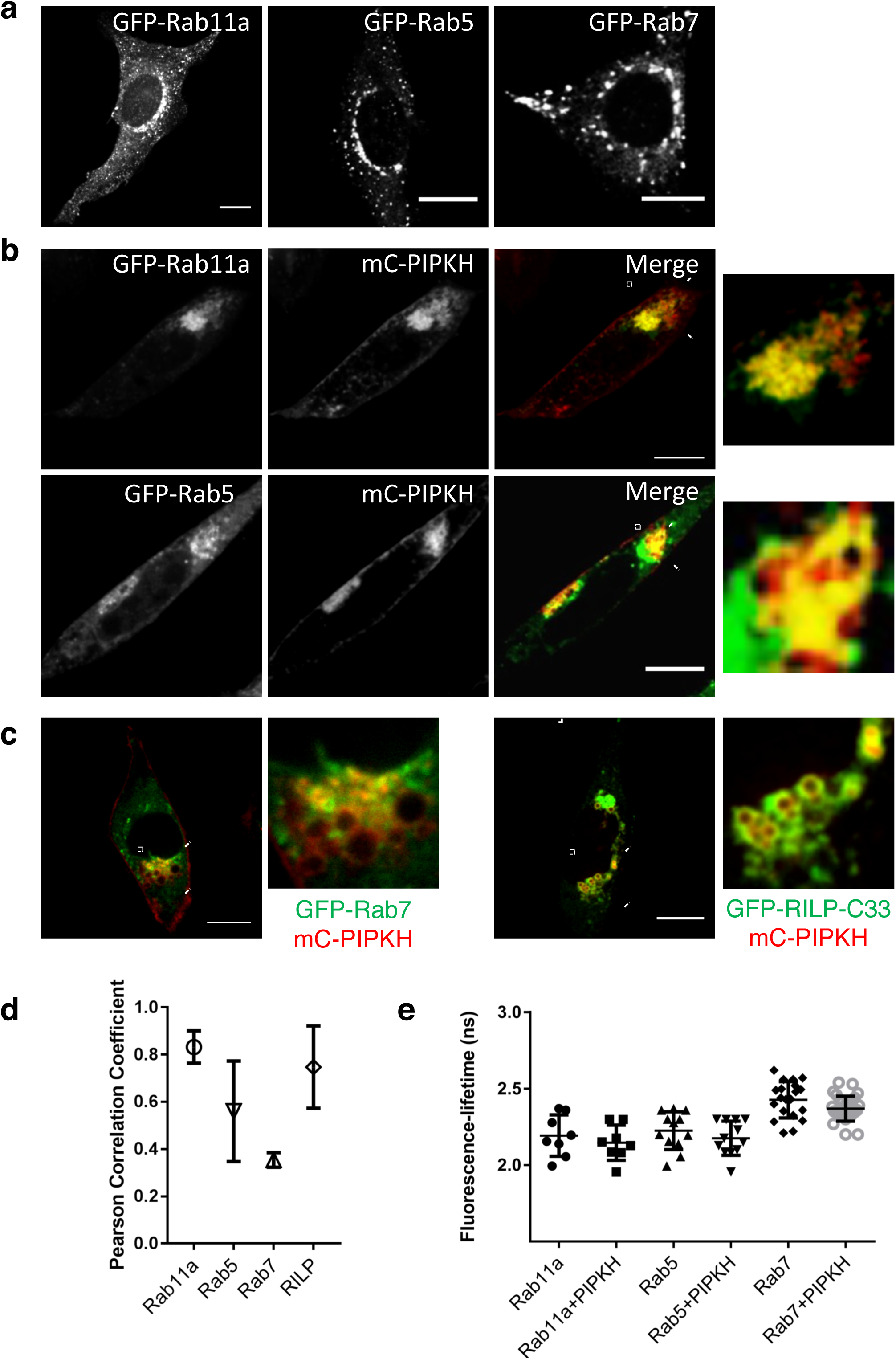
PIPKH colocalizes with endocytic Rab proteins. (a) Distribution of classical endocytic Rab proteins in CHO cells. CHO cells were transiently transfected with GFP tagged versions of Rab11a, Rab5 and Rab7 and imaged by confocal microscopy 18-24 h post-transfection. (b) PIPKH colocalizes with Rab11a and Rab5 on clustered endosomes. CHO cells were co-transfected with mCherry (mCh)-PIPKH and GFP chimeras of either Rab11a and Rab5 and images captured using spinning disc confocal microscopy. (c) PIPKH localizes with activated Rab7. CHO cells were co-transfected with mCh-PIPKH and either GFP chimeras of Rab7 or Rab7-interacting lysosomal protein (RILP) and imaged by spinning disc confocal microscopy. (d) Pearson’s correlation coefficient for the indicated Rab proteins, active Rab7 and PIPKH. Values represent the mean ± Std dev from a minimum of 9 cells per experiment from at least three different experiments (n=3). (e) PIPKH colocalizes with but is not proximal to the indicated Rab proteins. Distribution of fluorescence lifetimes of GFP (donor) tagged to the specific Rab protein and mCh-PIPKH. Values are the mean ± Std dev from a minimum of 9 technical replicates from at least three different experiments. Statistical testing was performed using unpaired t-test with Welch correction. Note: there is no significant difference in lifetime between donor alone and donor plus acceptor for these pairs of proteins. Scale bars = l0 mm.

**Figure 4:**
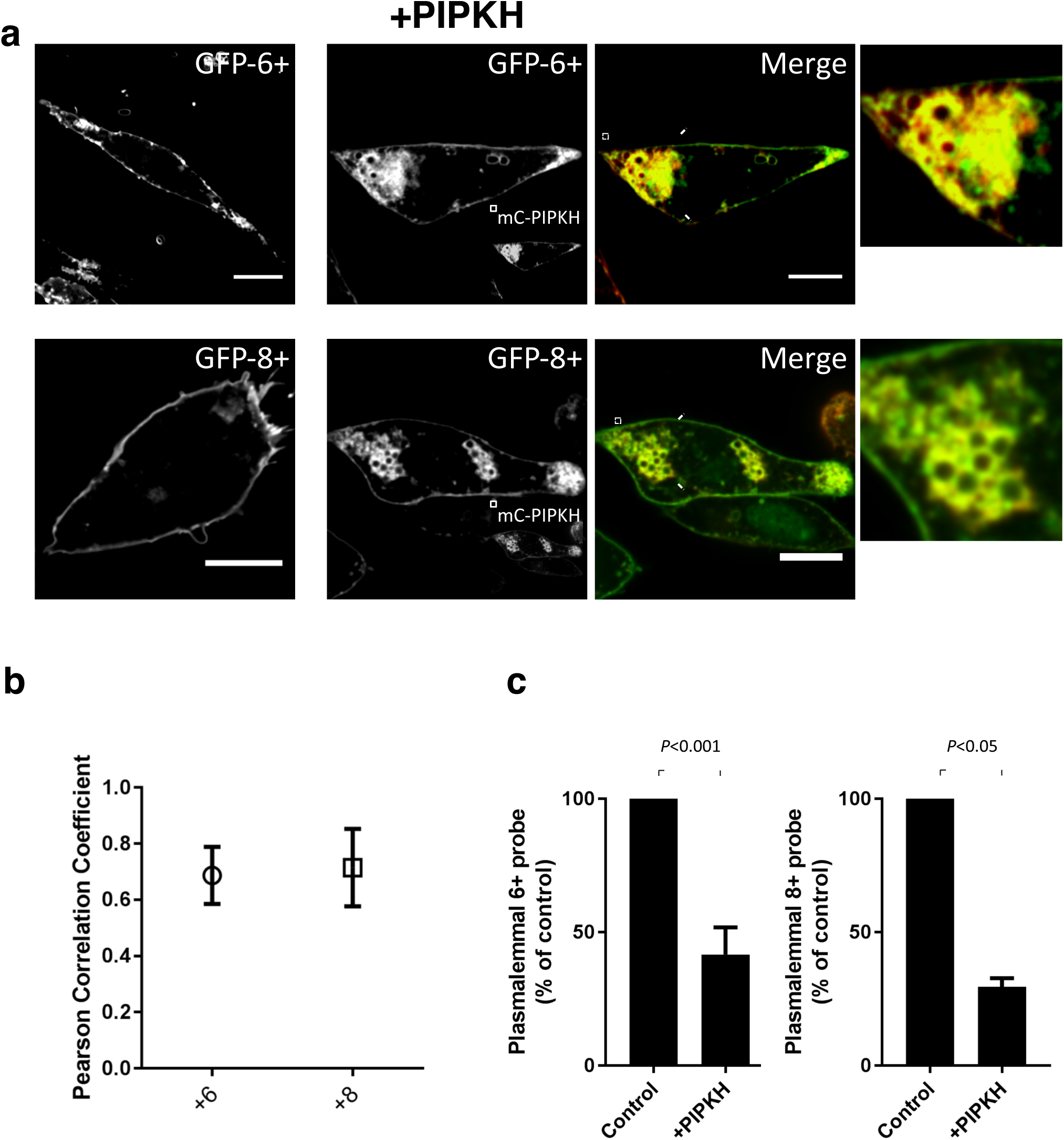
Enhanced expression of PIPKH alters the membrane surface charge of endosomal compartments. (a) Expression of PIPKH causes a redistribution of surface charge sensitive probes. CHO cells were transiently transfected with GFP tagged charge probes, +6 and +8, or were transiently co-transfected with mCherry ‐PIPKH and GFP-+6 or GFP-+8 charge probe. Images were captured using spinning-disc confocal microscopy l8-24 h post-transfection. (b) PIPKH colocalizes with the negative surface charge detectors. Pearson’s correlation coefficient plot for PIPKH and GFP-+6 and GFP-+8 is depicted. Values are the mean ± Std dev from a minimum of 9 technical replicates per day and at least three different experiments (n=3). (c) PIPKH stimulated alterations in phosphoinositide distribution cause a decrease in the amount of charge probe associated with the PM. Histogram represents the percent of charge probe associated with the PM in control and PIPKH-expressing cells. The percentages ± Std dev from a minimum of 9 cells from at three different experiments. Statistical testing was performed using a paired two-tailed Student’s t-test with Welch correction.

### PIPKH acts to relocalize polycationic proteins K-Ras and Rac1

We hypothesized that the PIPKH-induced aberrant PIP distribution would cause an increase in the negative charge density on endosomal compartments. To test this hypothesis, CHO cells were transfected or cotransfected with surface charge biosensors and PIPKH. As illustrated in Fig. 5a-c, CHO cells transfected with charge biosensors alone, both of the +8 and +6 charge biosensors localized preferentially to the plasma membranes consistent with previous findings^22^.

**Figure 5:**
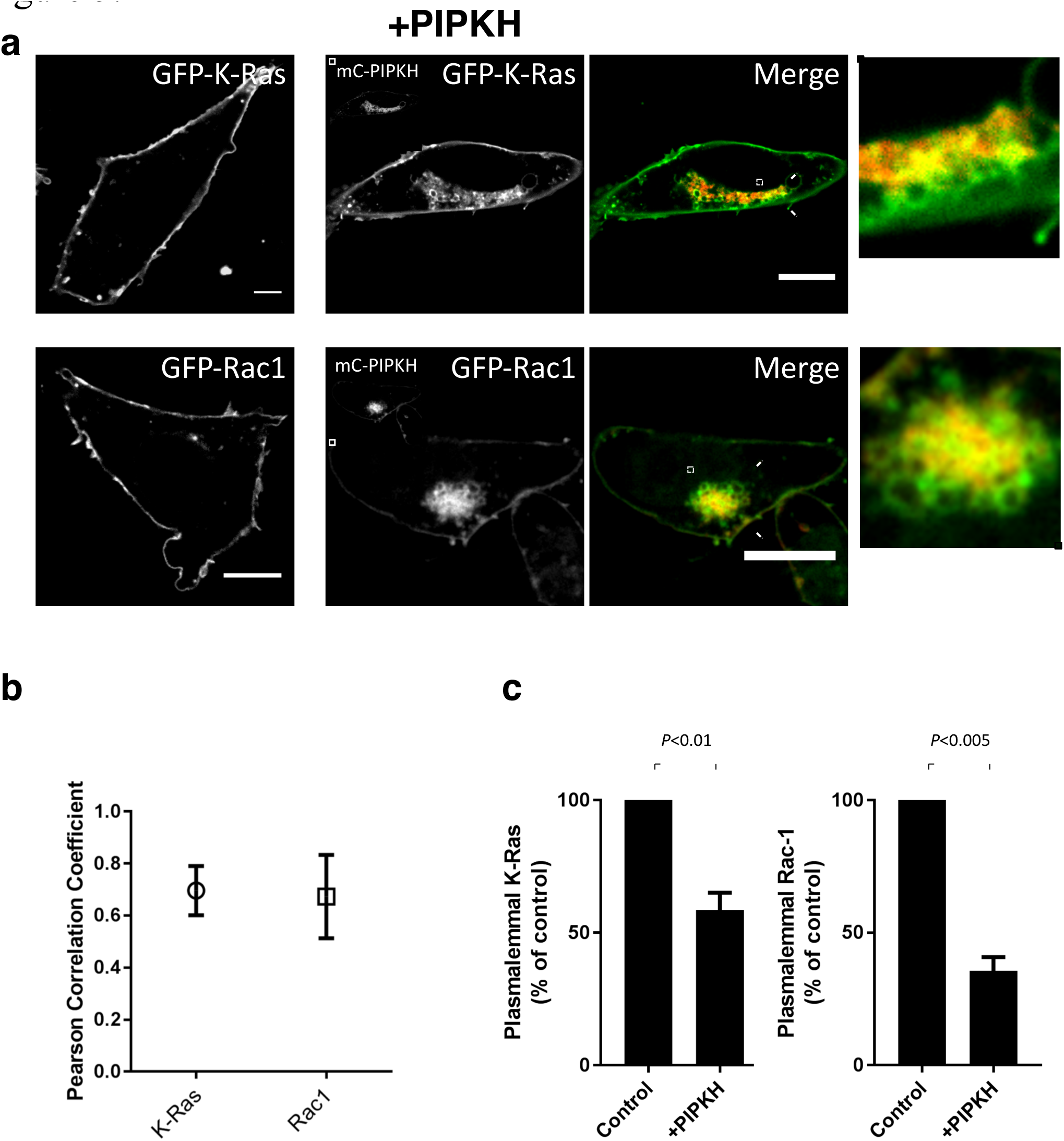
PIPKH overexpression relocalizes K-Ras and Rac1 to endosomal compartments. (a) Expression of PIPKH causes a redistribution of small G-proteins with polybasic tails. CHO cells were transiently transfected with GFP tagged versions of K-Ras and Rac1 or were transiently co-transfected with mCherry-PIPKH and GFP-Rac1 or GFP-K-Ras. Images were captured using spinning-disc confocal microscopy l8-24 h post-transfection. (b) PIPKH causes an accumulation of the small G-proteins on endosomes. Pearson’s correlation coefficient plot for PIPKH and GFP-K-Ras and GFP-Rac1 are plotted. Values are the mean ± Std dev from a minimum of 9 technical replicates and at least three different experiments (n=3). (c) PIPKH stimulated alterations in phosphoinositide distribution cause a decrease in the amount of K-Ras and Rac1 associated with the PM. Histogram represent the percent of the indicated small G-protein associated with the PM in control and PIPKH-expressing cells. The percentages ± Std dev from a minimum of 9 technical replicates per biological replicate with n=3. Statistical testing was performed using a paired two-tailed Student’s t-test with Welch correction.

Conversely, in CHO cells transiently expressing PIPKH, the surface charge biosensors were co-localized with PIPKH to endosomal compartments suggesting that significant phosphoinositide is being generated to impact the charge probes. Next, we extended this finding to small GTPases that use polycationic patches to localize to the PM, K-Ras and Rac1. To do this we transiently co-transfected K-Ras or Rac1 with PIPKH in CHO cells. Compared to the control cells, both K-Ras and Rac-1 were observed to relocalized from the PM to PIPKH-positive endosomal compartments (Fig. 6a-c). These results indicate that PIPKH-induced alterations in the cellular distribution of PIPs is sufficient to cause relocalization of proteins that normally target to the PM via electrostatic interactions. These results support the observations that the degree of negative surface charge on a membrane can dictate the association of peripheral membrane proteins.

## Discussion

In this paper, we have assessed the ability of the pseudokinase PIPKH to influence PIP5Ks and thus the production of PtdIns(4,5)P_2_ within the cell. Cells overexpressing mCh-PIPKH showed different localization patterns for the PIP5Ks. Consistent with a previous report that the impact of PIPKH on PIP5Ks appears to be direct protein-protein interactions^18^. We found that the fluorescence lifetime of the GFP/YFP-tagged versions of PIP5Ks was decreased in the presence of mCh-PIPKH. Together these results suggest that PIPKH can interact with PIP5Ks and thereby regulating their cellular localization. Intriguingly, the relocalization of PIP5Ks and/or the accumulation of endosomal PtdIns(4,5)P_2_ was associated with the clustering of endosomes in a perinuclear region. However, the mechanisms driving this apparent clustering is unclear. One possibility is that peripheral proteins associated with the limiting membrane of the endosomes may be altered following PIPKH expression. Indeed, the polyvalent anionic phosphoinositides contribute to confer a negative surface charge on the inner leaflet of the plasma membrane, which is critical for the targeting of many plasmalemmal proteins^7,8^. In support of this notion, we find that changes in cellular PtdIns(4,5)P_2_ distribution causes a mislocalization of plasmalemmal proteins to PIPKH-/PtdIns(4,5)P_2_-positive endosomes. Together, our results suggest that the endogenous function of PIPKH may include acting as a regulatory protein for PIP5Ks to modulate the abundance of PtdIns(4,5)P_2_ in the particular compartment within the endocytic pathway.

The specific regulation of PIP5K activity and cellular localization is essential for generating cellular pool or local concentrations of PtdIns(4,5)P_2_. It is well known that a 25-amino acid segment, named the activation loop, confers PIP5Ks substrate specificity to PtdIns4P^23^. PIP5Ks colocalize with their product PtdIns(4,5)P_2_, in the PM in a variety of cell types including macrophages by virtue of a positively charged face^6^. Several proteins have been described to influence the activity of PIP5K isoforms via direct interactions^24–26^. This includes small GTPases Rho, Rac and ARF that have been describe to aid in the targeting of PIP5Ks *in vivo* and stimulate the kinase activity *in* vitro^15–17,26^. These interactions may help facilitate the generation of specific pools of PtdIns(4,5)P_2_. For instance, Rac has been demonstrated to influence PIP5Kβ at the PM during neurite retraction^27^. While the PH-PLC8 typically only decorates the PM, PIP5Ks and their product have also been described to be present in a variety of other, more difficult to observe, subcellular locations including, endosomes, autophagolysosomes and the nucleus^28–33^. We suspect that many of these specific pools will be controlled by proteinprotein interactions.

PIPKH likely arose from gene duplication of one of the functional PIPK5Ks but has been rendered catalytically inactive by mutations^18^. The maintenance of the intact PIPKH open reading frame (ORF) in fish, reptiles, birds and mammals vertebrates–roughly 400 million years of evolution–argues against the gene being a pseudogene instead of a pseudokinase^34^. Unfortunately, the endogenous function of PIPKH remains unclear in mammals. Proteomics and expression data suggest that PIPKH is present in gastric epithelial cells^19^. While PIPKH has been characterized as being lost in a subset of gastric cancer samples that correlated with enhanced cell proliferation and migration^20^. However, further work towards improving understanding of the biophysical properties of PIPKH is required for revealing its precise role(s) in cancer metastasis. It is known that gastric epithelial cells express caveolin and possess caveolae. In this regard, the only other study that we are aware of that has directly investigated the cellular impact of PIPKH was an siRNA-mediated screen of the human kinome and the impact on caveolae. In this study, Pelkams and Zerial discovered that silencing of PIPKH in HeLa cells to an increase of multi-caveolar assemblies or "rosettes" at the expense of individual caveola^23^. The main caveat of this study was that the efficiency and specificity of the silencing was not investigated. Additionally, it is not well understood what signals control the formation of higher ordered caveolar assemblies compared to the traditional standalone caveola.

In summary, our results indicate that PIPKH may function as a regulatory protein for PIP5Ks. Based on its localization we suspect that PIPKH supports the production of a specialized pool of PtdIns(4,5)P_2_ in the endocytic pathway. Collectively, our results together with previous work suggest that PIPKH may only be important in a specialized cell type such as the gastric epithelial cells. However, due to its preservation over a long evolutionary time scale an alternative possibility is that it supports a key developmental process not present in immortalized cultured cells. This will require further investigation in the future.

## Methods

### Plasmids

Full length human PIPKH was synthesized by Integrated DNA Technologies (Coralville, IA, USA) and subcloned into the pIDT-Amp plasmid. PIPKH was amplified by PCR using this plasmid as a template using the following pairs of primers: 5’-CGCGGATCCTCACTCTGTATGGGCTTCTA-3’ and 5’-GCGGTCGACATGGCTGCACCATCACCAGG-3’. The PCR product was introduced into pmCherry-C1 and pEGFP-C1 vector using the restriction enzymes SalI and BamHI site. The GFP/YFP-PIP5Ks were published previously^6^. The following plasmids used in this study have been previously described: GFP-P4Mx2^35^, mCh-p40-PX^36^, GFP-PH-PLC8 and GFP-AKT-PH^37^, GFP-Rab5^38^, GFP-Rab7 and GFP Rab11a^39^, GFP-EEA1^40^, GFP-RILP-C33^21^. The surface charge probes and small G-proteins, GFP-6+, GFP-8+, GFP-K-Ras, and GFP-Rac1, were kind gifts of Dr. Sergio Grinstein (The Hospital for Sick Children, Toronto, ON, CAN)^22^.

### Cell culture and transfection

Chinese hamster ovary (CHO) cells were maintained at 37°C with 5% CO2 in Ham’s F12 and supplemented with 10% fetal bovine serum from Wisent (Burlington, ON, CAN). CHO cells were transiently transfected with plasmids using Fugene 6 (Promega, Madison, WI, USA) according to the manufacturer’s instructions. The following day, 18-24 h post-transfection, cells were observed or fixed with 3.7% paraformaldehyde (Electron Microscopy Sciences, Hatfield, PA, USA) in phosphate-buffered saline for 60 min at room temperature, washed with Tris-buffered saline and stored at 4 °C. Fixed samples were typically imaged within 48 h of fixation.

### Microscopy

#### Confocal microscopy

Fluorescence images were acquired using spinning-disc confocal microscopy or as indicated laser scanning microscopy. The spinning-disc confocal system used is located within the Imaging Facility at the Hospital for Sick Children, Toronto, Ontario. This system assembled by Quorum Technologies (Guelph, Ontario) and is based on an Olympus IX81 with a 60×/1.35 NA oil immersion objective. The system is equipped with diode-pumped solid-state laser lines (440, 491, 561, 638, and 655 nm; Spectral Applied Research), motorized XY stage (Applied Scientific Instrumentation), and a piezo focus drive (Improvision). Images were acquired using back-thinned, electron-multiplied cooled charge-coupled device (EM-CCD) cameras (Hamamatsu Photonics) driven by the Volocity software (version 6.3.0; PerkinElmer).

The laser scanning microscope used is located in the St. Michael’s Hospital BioImaging Facility is a Zeiss LSM 700 inverted confocal microscope with a Plan-Apochromat 60x/1.4 NA oil objective and Zen 2010 software (Zeiss). Analysis of images was performed using Volocity, Zen 2010 or ImageJ software (NIH, Bethesda, MD, USA) depending on the experiment.

#### Fluorescence lifetime measurement - fluorescence resonance energy transfer (FLIM-FRET)

FLIM-FRET measurements were performed at the Hospital for Sick Children Imaging Facility using an Olympus IX81 equipped with light emitting diodes LED lines (402, 446, 483, and 540 nm) and a Lambert Instruments FLIM (LI-FLIM) Attachment. Images were acquired using an 150x/1.45 NA oil immersion objective and a Hamamatsu C9100-13 Electron Multiplying (EM)-CCD camera using the 483 and 540 LED excitation lines. The LI-FLIM v1.2.12 (Lambert Instruments, Groningen, Netherlands, NLD) was used to monitor the fluorescence lifetimes of the donor fluorophores.

### Post-acquisition analysis

The degree of colocalization was determined by Pearson’s Correlation Coefficient (or simply Pearson’s) using the JACoP plugin in ImageJ. Specifically, cells were selected using the freehand tool in ImageJ. Images were then split into the two contributing channels for further analysis. For consistent and reproducible analysis, auto-thresholding for the two channels was performed using the Yen algorithm as previously described followed by the determination of the Pearson’s^41^.

For determining the relative distribution of fluorescence probes/proteins in the cell the ROI tool was used to highlight areas or the PM, cytoplasm or background (outside of the cell) and determine the mean intensity/pixel. The percent of plasmalemmal resident probe was determined by subtracting the integrated cytoplasmic signal from the total cellular fluorescence and represented as a percent of the total cellular fluorescence.

## Acknowledgements

We thank Mr. Paul Paroutis and Ms. Kimberly Lau (Hospital for Sick Children) and Dr. Caterina Di Ciano-Oliveira (St. Michael’s Hospital) for expert assistance with microscopy and image analysis. This work is supported by a Natural Sciences and Engineering Research Council (NSERC) Discovery Grant to G.D.F. G.D.F. is a recipient of a New Investigator Award from Canadian Institutes of Health Research (CIHR); and an Early Researcher Award from the Government of Ontario. Y.Y. has been supported in part by a scholarship from St. Michael’s Hospital and a scholarship from the Alzheimer Society of Canada. M.P. was a recipient of an NSERC undergraduate student researcher award (USRA).

## Author contributions statement

G.D.F. conceived the experiment, oversaw the project, and wrote the manuscript. Y.Y. conducted the experiments, analysed the results and wrote the first draft of the manuscript. M.P. generated novel reagents and conducted preliminary experiments. All authors reviewed the manuscript.

## Additional information

The authors declare no competing or financial interests.

## Figure Legends

**Supplemental Figure 1.**
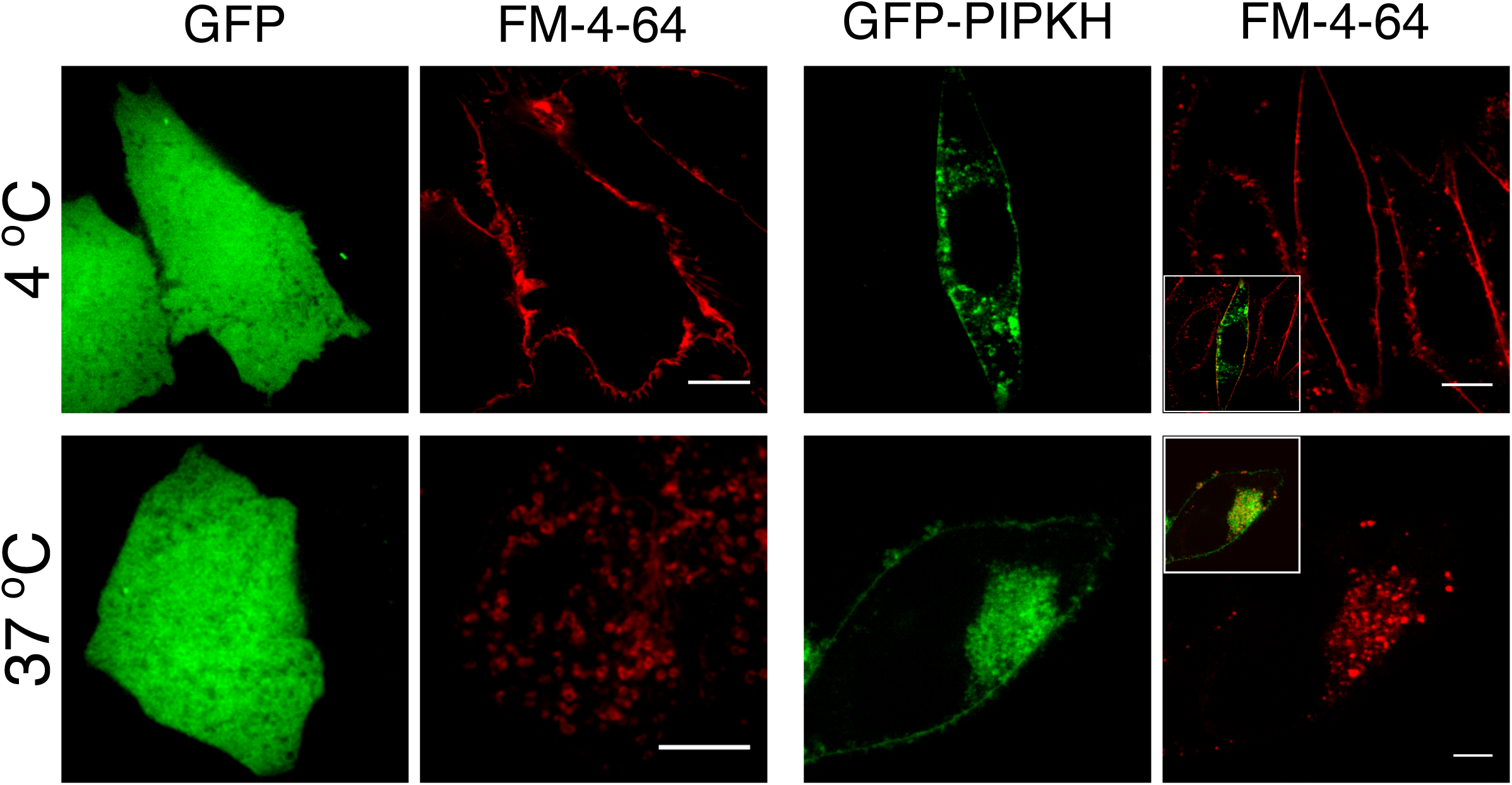
Cytoplasmic PIPKH structures are endocytic and are not connected to the plasma membrane. (a, b) PIPKH localizes to endocytic structures. CHO cells transiently expressing GFP-PIPKH were stained with FM4-64 for 30 min at either 4 °C (a) or 37 °C (b) for 30 min. Depicted are representative images. FM4-64 delineates the plasma membrane but does not label the cytoplasmic PIPKH positive structures, demonstrating that these are not invaginations that remain connected with the plasma membrane. Following a 30 min continuous pulse with FM4-64, the majority of PIPKH positive structures contain the endocytic marker. Scale bars = 10 mm.

